# Chromosome-level genome assembly and annotation of the chaetognath *Flaccisagitta enflata*

**DOI:** 10.1101/2025.03.14.643333

**Authors:** Nadège Guiglielmoni, Michael Eitel, Pierrick Moreau, Stefan Krebs, Mark Vermeij, Romain Koszul, Jean-François Flot

## Abstract

Chaetognaths, a phylum of enigmatic marine predators, present a significant challenge to phylogenetic reconstruction due to their uncertain evolutionary placement. While transcriptome analyses have suggested affinities with the Gnathifera clade, genomic data for this group remain scarce, hindering a comprehensive understanding of their evolution. Here, we present the first chromosome-level genome assembly of Flaccisagitta enflata, a species within the Aphragmophora order. The genome assembly includes 9 chromosome candidates with a total size of 794 Mb and a BUSCO score of 91.3% against the Metazoa lineage. This high-quality genome assembly provides a crucial resource for comparative genomic analyses within Chaetognatha and the broader Gnathifera clade, and it will facilitate investigations into chaetognath evolution and their phylogenetic relationships, addressing long-standing questions regarding their placement within the animal kingdom.

## Background

Chaetognaths, or arrow worms, are transparent marine predators found at various depths across all oceans, though most species prefer shallow waters [1]. They have an elongated, transparent body with one or two pairs of lateral fins, a caudal fin, a distinctive head equipped with hooks, and their size ranges from a few millimeters to several centimeters [2]. Chaetognaths are classified into two orders based on the presence of transverse muscles, or phragms: Phragmophora and Aphragmophora [3]. The phylum comprises approximately 150 known species and represents an enigmatic clade with an uncertain phylogenetic position. Initially, chaetognaths were thought to be deuterostomes due to their developmental characteristics, but 18S rDNA analyses disproved this hypothesis [4, 5] and further supported the division into Phragmophora and Aphragmophora [6]. Later, Nielsen proposed that chaetognaths belong to the Gnathifera clade, which includes Gnathostomulida, Micrognathozoa, and Rotifera [7]. This hypothesis has recently been reinforced by transcriptome analysis of ten chaetognath species [8], yet no high-quality genome assembly was generated so far for this group due to the challenge of sequencing and assembling non-model species [9], and only one chromosome-level genome assembly of the rotifer *Adineta vaga* is available for Gnathifera [10].

We present a first and chromosome-level genome assembly of a species belonging to the phylum Chaetognatha, namely *Flaccisagitta enflata*. This species was initially described by Grassi in 1881, and is classified in the order Aphragmophora and the family Sagittidae. This species is found at low depth and in seas at more elevated temperatures [11] and has some large specimens reaching up to 2.5 cm. Its genome size is moderate, estimated at 0.71 pg using Feulgen Image Analysis Densitometry [12].

## Methods

### Sample collection and fixation

Chaetognaths were collected in Snake bay, Curacao, on November 2nd 2019 after sunset. Individuals were identified as *Flaccisaggita enflata* following the key provided in [11] and based on the following criteria: length up to 2.5 cm; body transparent, soft and inflated-looking; 8-10 hooks (Figure 1.A1); no collarette; anterior position of the ganglion (Figure 1.A2); pair of lateral fins; short rounded fins; ovaries not extending till the anterior fins (Figure 1.B3); round vesicles close to the tail (Figure 1.B4).

**Figure 1:**
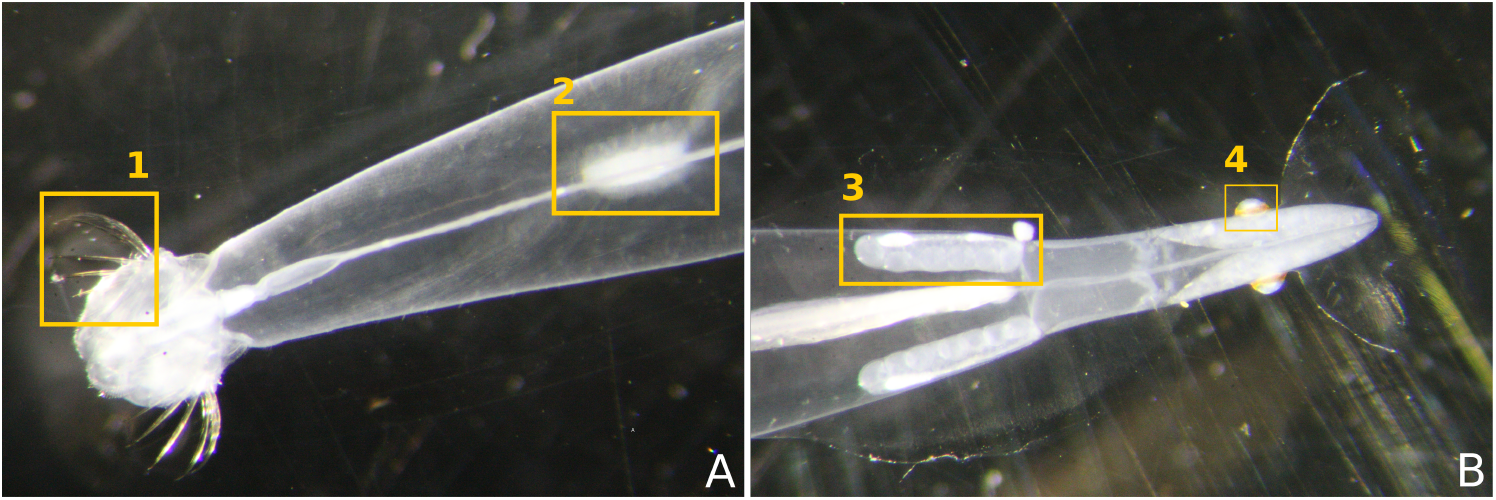
Crosslinked individual C2 used for Hi-C sequencing.

One 2-cm individual C1 was preserved in 96% ethanol and kept at 4°C for high-molecular-weight DNA extraction. One 1.5-cm individual C2 was crosslinked in 3% formaldehyde for 45 minutes, quenched in 250 mM glycine, then frozen at -80°C for Hi-C. Two individuals C3 and C4 of 1.5 and 1.2 cm were preserved in RNAlater and kept at 4°C for RNA extraction.

### High-molecular-weight DNA extraction and sequencing

C1 was extracted following a modified version of the protocol described in [13]. The whole individual was incubated in 180 µL of cetyltrimethylammonium bromide (CTAB) buffer (polyvinylpyrrolidone 2%, tris(hydroxymethyl)aminometha HCl 100 mM, ethylenediaminetetraacetic acid 25 mM, NaCl 2 M, CTAB 2%, *β*-mercaptoethanol 1/100) and 25 µL of proteinase K for 3 hours at 60°C and 300 rpm. The lysed sample was purified with phenol-chloroform-isoamyl alcohol 25:24:1, chloroform-isoamyl alcohol 24:1 and with AMPure XP beads, which yielded 2.5 µg of DNA with OD260/280=1.95 and OD260/230=1.95.

A long-read library was prepared for C1 with the Nanopore SQK-LSK109 Ligation sequencing kit and sequenced using a PromethION flow cell (with pore proteins R9.4.1), which ran for 72 hours with 3 nuclease flushes and reloads, and gave an output of 40.5 Gb with an N50=6.1 kb. Basecalling was done with Guppy v4. 168 Gb of paired-end 150-bp Illumina reads were additionally sequenced for C1 by Novogene on a NovaSeq.

### Hi-C sequencing

The crosslinked individual C2 was used to prepare a Hi-C library with the Arima Hi-C kit (including the restriction enzymes DpnII and HinfI), and resulted in 466 ng of DNA. The DNA was fragmented with a Covaris (300 bp) and biotinylated fragments were selected with streptavidin beads. The library was prepared for Illumina sequencing using the Invitrogen TM Collibri TM PS DNA Library Prep Kit and following manufacturer instructions. Sequencing by Novogene resulted in 489 millions pairs of 150-bp reads.

### RNA extraction and sequencing

RNA was extracted by Novogene using a Trizol-based protocol. Samples were sequenced on a NovaSeq platform and yielded 320 and 122 millions pairs of 150-bp reads for C3 and C4 respectively.

### Pre-assembly analysis

The genome size was estimated with BBtools v38.79 [14] using the script `kmercountexact.sh` and the shotgun Illumina reads. As this tool is based on *k* -mers, three values of *k* were tested with *k* = *{*27, 29, 31*}*. A *k* -mer histogram of the Illumina dataset was built using KAT hist v2.4.2 [15] (with *k* =27).

### Genome assembly

Shotgun Illumina and Hi-C reads were trimmed using cutadapt v2.9 with parameter `-m 10` [16]. Nanopore reads were trimmed using Porechop v0.2.4 [17] with default parameters, corrected using the Illumina reads and Ratatosk v0.7.5 [18] with default parameters, and assembled *de novo* using Canu v1.9 [19] with parameter `genomeSize=700m` and the reads passed with `--nanopore-corrected`. Nanopore reads were mapped to the contigs using minimap2 v2.17 [20] with parameter `-x map-ont` and uncollapsed haplotypes were purged with purge dups [21] twice with default parameters. Hi-C reads were mapped to the draft assembly using hicstuff v2.3.0 [22] and bowtie2 v2.3.5.1 [23] with parameters `--enzyme DpnII,HinfI --iterative`. instaGRAAL v0.1.6 no-opengl branch [24] was run with the parameters `--level 5 --cycles 150`. The scaffolds were post-processed with `instaGRAAL-polish` to reduce misassemblies and 10 Ns were added as gaps. Gaps in the chromosome candidates were filled using TGS-GapCloser [25] and the Ratatosk-corrected Nanopore reads with parameter `--ne`, and the scaffolds were further polished using HyPo [26] with parameters `-c 240 -s 700` after mapping the Illumina reads with bowtie2.

### Assembly evaluation

Assemblies were assessed using BUSCO v5.8.0 [27] against the lineage Metazoa odb10 with parameter `-m genome` and using KAT comp v2.4.2 [15] against the Illumina dataset (with *k* =27) with default parameters. The Hi-C contact map was built using the tool hicstuff with `hicstuff pipeline` as described previously and `hicstuff view` with the parameter `--binning 2000`.

### Genome annotation

Transposable elements (TE) were annotated with the EDTA pipeline v1.9.8 [28] with parameters `--sensitive 1 --anno 1 --overwrite 1 --force 1`: first, long-terminal repeat (LTR) candidates detected by LTRharvest v1.6.1 [29] and LTR FINDER v1.07 [30] were filtered with LTR retriever v2.9.0 [31]; Helitron transposons were searched for with HelitronScanner [32], while other repeats were detected by Generic Repeat Finder v1.0 [33] and TIR-learner [34]; a final TE library was produced after further filtering and repetition annotation with RepeatModeler v2.0.1 [35]. The output hardmasked assembly was converted into a softmasked assembly. RNA-seq reads were trimmed with cutadapt v2.9 [16] with parameter `-m 10`. C3 reads were assembled with Trinity v2.12.0 [36]. This transcriptome and the C3 and C4 reads were provided as input to Funannotate [37]. `funannotate train` was run with parameters `--no trimmomatic --no normalize reads` to map the transcripts to the softmasked assembly using minimap2 v2.17 [20] and produce a first annotation with PASA v2.4.1 [38]. These files were used by `funannotate predict` for gene prediction with GeneMark-ES [39] and Augustus [40], and a consensus gene set was produced by Evidence Modeler [41]. Annotation was evaluated using BUSCO v5.8.0 [27] against the lineage Metazoa odb10 with parameter `-m proteins`.

### Genome quality

The full assembly pipeline is presented in Figure 2a. Genome size was estimated to 695-699 Mb (haploid) with BBTools, matching the cytological estimation of 0.71 pg. Initial contigs were obtained by assembling Ratatosk-corrected Nanopore reads with Canu and reached 1.48 Gb, close to the expected size for a diploid genome assembly, with an N50 of 195 kb. Contigs were purged using purge dups and resulted in an assembly of 946 Mb with an N50 of 215 kb. The assembly was scaffolded using instaGRAAL and yielded 9 chromosome candidates ranging from 71.5 to 112.6 Mb. Gaps were filled using TGS-GapCloser and the final scaffolds were polished using HyPo, to obtain a final assembly of 794 Mb with an overall BUSCO score of 91.3% against the Metazoa odb10 lineage (Table 1). Sequences not anchored to the main scaffolds were removed from the assembly to prevent contamination, which is expected as the samples were preserved directly after collection and not starved nor cleaned. Plankton on which chaetognaths feed would hardly be distinguishable from the host using typical decontamination strategies, as they have linear genomes, a GC content similar to chaetognaths, and are likely poorly documented in public databases. The *k* -mer plot (Figure 2b) shows two peaks, as is expected for a diploid species. Erroneous *k* -mers at low multiplicity are not included in the assembly (0X, in black). Limited artefactual duplications are shown in the homozygous peak (2X, purple). The 9 chromosome candidates can be identified on the contact map (Figure 2c) based on their heightened intrachromosomal contacts. 64.2% of the assembly was identified as repetitive and 35,833 genes were predicted with a BUSCO score of 87.8% (77.0% single-copy, 10.8% duplicated features) against the Metazoa odb10 lineage.

**Table 1:** Assembly statistics.

|  | **Final scaffolds** |
| --- | --- |
| Assembly size | 794 Mb |
| N50 | 91.2 Mb |
| # fragments | 9 |
| Gaps | 1,476 |
| Ns | 23,555 |
| Single-copy BUSCO | 80.2% |
| Duplicated BUSCO | 11.1% |

**Figure 2:**
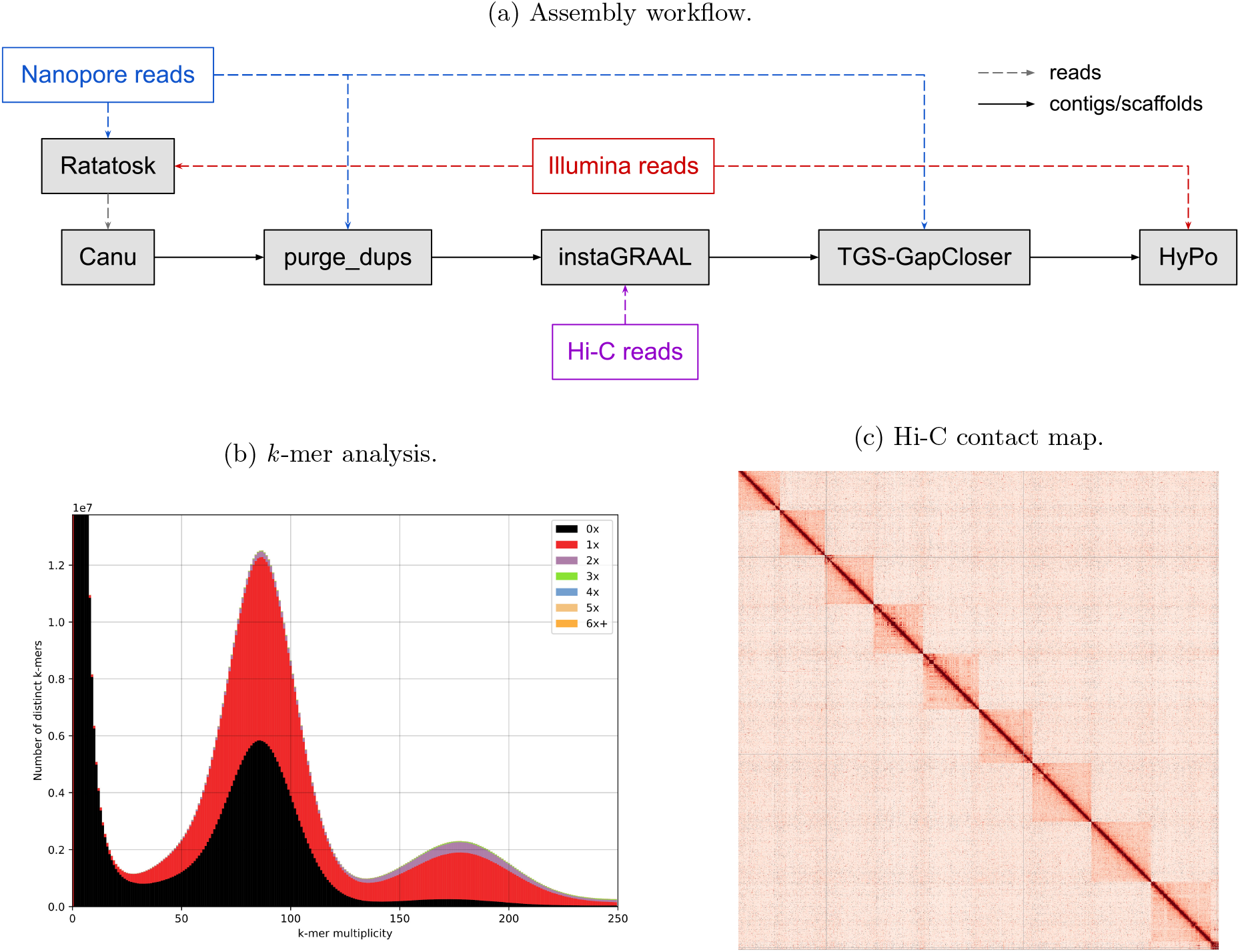
a) Assembly workflow. Nanopore reads were corrected using Ratatosk with Illumina reads. Both Nanopore and Illumina reads were generated from a DNA extraction of individual C1. Corrected reads were assembled using Canu and haplotigs were removed using purge dups. The assembly was scaffolded using instaGRAAL and Hi-C reads from individual C2. The scaffolds were gap filled using TGS-GapCloser and the Nanopore reads, and polished using HyPo and the Illumina reads. b) *k* -mer analysis using KAT comp with *k* =27. The spectrum shows two peaks, the first one for heterozygous *k* -mers and the second one for homozygous *k* -mers. c) Hi-C contact map of the final scaffolds. 9 chromosome candidates with heightened intrachromosomal contacts can be identified.

### Perspectives

The chromosome-level assembly of *Flaccisagitta enflata* provides a first reference for the phylum Chaetognatha, and shall be employed in comparative and phylogenomic studies. This genome assembly posed a challenge due to the high heterozygosity between haplotypes of the species, and accordingly we strived to use few individuals. Nanopore and shotgun Illumina reads were obtained from a single individual and can be used to generate a draft phased assembly. With the improvements of long-read sequencing technologies, it would be possible to obtain both long reads and Hi-C reads from a single individual of similar size by using a portion for low-input PacBio HiFi sequencing and the rest for Hi-C sequencing. The high-accuracy of PacBio HiFi reads and Hi-C would be an adequate combination to produce a haplotype-resolved assembly. A phased assembly would be most appropriate for this species due to its high heterozygosity and the difficulty of collapsing divergent regions [42]. DNA extraction may additionally be optimized by flash-freezing and crushing the sample prior to chemical lysis, and incubating the powder in CTAB buffer and proteinase K for 30 minutes to 1 hour maximum, as prolonged lysis in CTAB may explain suboptimal fragment length.

## Supplementary information

Command lines are listed in github.com/nadegeguiglielmoni/Flaccisagitta enflata methods.

## Data availability

Sequencing datasets and assembly were submitted to European Nucleotide Archive under the accession number PRJEB83807.

## Acknowledgements

We thank Christophe Chapard for his help in preparing the Hi-C library.

